# Establishing Synthesis Pathway-Host Compatibility via Enzyme Solubility

**DOI:** 10.1101/386144

**Authors:** Sara A. Amin, Venkatesh Endalur Gopinarayanan, Nikhil U. Nair, Soha Hassoun

## Abstract

Current pathway synthesis tools identify possible pathways that can be added to a host to produce a desired target molecule through the exploration of abstract metabolic and reaction network space. However, not many of these tools do explore gene-level information required to physically realize the identified synthesis pathways, and none explore enzyme-host compatibility. Developing tools that address this disconnect between abstract reactions/metabolic design space and physical genetic sequence design space will enable expedited experimental efforts that avoid exploring unprofitable synthesis pathways. This work describes a workflow, termed Probabilistic Pathway Assembly with Solubility Scores (*ProPASS*), which links synthesis pathway construction with the exploration of the physical design space as imposed by the availability of enzymes with characterized activities within the host. Predicted protein solubility propensity scores are used as a confidence level to quantify the compatibility of each pathway enzyme with the host (*E. coli*). This work also presents a database, termed Protein Solubility Database (*ProSol DB*), which provides solubility confidence scores in *E. coli* for 240,016 characterized enzymes obtained from *UniProtKB/Swiss-Prot*. The utility of *ProPASS* is demonstrated by generating genetic implementations of heterologous synthesis pathways in *E. coli* that target several commercially useful biomolecules.

**Availability:** *ProSol DB* data and code for *ProPASS* are available for download from https://github.com/HassounLab/

## Introduction

Synthesis pathways have been engineered within microbial hosts to produce commercially useful biomolecules including polyesters (Fidler & Dennis, 1992), building blocks for industrial polymers (Tong, Liao, & Cameron, 1991), biofuels (Nawabi, Bauer, Kyrpides, & Lykidis, 2011; Radakovits, Jinkerson, Darzins, & Posewitz, 2010; E. J. Steen et al., 2010), and therapeutic natural products derived from isoprenoids (Martin, Pitera, Withers, Newman, & Keasling, 2003; Pitera, Paddon, Newman, & Keasling, 2007; Watts, Mijts, & Schmidt-Dannert, 2005), polyketides (Peirú, Menzella, Rodríguez, Carney, & Gramajo, 2005; Pfeifer, Admiraal, Gramajo, Cane, & Khosla, 2001), and non-ribosomal peptides (Takahashi et al., 2007). Synthesis pathways consist of series of non-native enzyme-controlled reactions from metabolites within the host to a target molecule. Often, many possible biochemical routes to a target molecule exist such as in the production of 3- hydroxypropionic acid through glycerol, β-alanine, or acrolein pathways (Cheng, Jiang, Wu, Li, & Ye, 2016; Luo et al., 2016; Song, Kim, Cho, & Lee, 2016). Further, the genetic implementation of each pathway is not unique since genes can be sourced from various organisms. For example, production of thebaine and hydrocodone in *S. cerevisiae* required construction of a heterologous pathway with enzymes from a bacterium (*Pseudomonas putida*), a mammal (*Rattus norvegicus*), and several plants (*Papaver somniferum*, *Papaver bracteatum*, *Coptis japonica*, *Eschscholzia californica*) (Galanie, Thodey, Trenchard, Interrante, & Smolke, 2015).

Needless to state, experimental efforts to explore all possible pathways and all possible organism-specific enzyme selections are prohibitive. Unfortunately, current pathway synthesis computational tools provide solutions based on exploring the abstract metabolic space without regards to possible genetic implementation. Identifying best genetic implementations are typically ad hoc and there is a lack of systematic design tools and workflows that support the co-exploration of the metabolic and genetic implementation design spaces. Pathway synthesis tools that consider genetic compatibility between biosynthetic pathways and heterologous hosts can significantly expedite the metabolic engineering cycle by focusing on design spaces with a greater fraction of profitable options.

In this light, *pathway synthesis tools* must conceptually perform two distinct tasks. The first task involves identifying possible synthesis pathways from the host to a target molecule and evaluating potential yield. This is known as the *pathway construction* or *pathway identification* problem, where sequences of reactions that synthesize the target molecule from a host metabolite are selected based on data from multi-organism databases such as KEGG (Kanehisa & Goto, 2000), MetaCyc (Caspi et al., 2008), and SEED (Overbeek et al., 2005). Pathway construction utilizes either rule-based techniques (e.g. BNICE (Wu, Wang, Assary, Broadbelt, & Krilov, 2011), PathPred (Moriya et al., 2010), Meta (Klopman, Dimayuga, & Talafous, 1994; Klopman, Tu, & Talafous, 1997), Meteor (Greene, Judson, Langowski, & Marchant, 1999; Marchant, Briggs, & Long, 2008), and UM-PPS (Ellis & Wackett, 2012; Hou, Wackett, & Ellis, 2003)) or graph-based techniques such as *ProPath* (Yousofshahi, Lee, & Hassoun, 2011), DESHARKY (Rodrigo, Carrera, Prather, & Jaramillo, 2008), Metaroute (Blum & Kohlbacher, 2008). Thus, there are currently multiple tools available for pathway construction.

The second synthesis task concerns the *parts selection problem* or, more broadly, the *pathway implementation* problem. This task involves identifying gene sequences from specific organisms that best realize reactions along the synthesis pathway. There are several factors to consider. One set of factors is associated with transcription and translation, including promoter strength and efficiency of ribosome binding. Another set of factors is dependent on the interactions of the enzyme’s protein coding sequence with the cellular metabolic machinery to ensure correct conformation and function while avoiding misfolding or aggregation, known as protein solubility. Overcoming protein solubility issues can be achieved by codon optimization, co-expressing molecular chaperones (Trésaugues et al., 2004; Xia et al., 2016), lowering culture temperature (Makrides, 1996) or modifying growth media (Makrides, 1996). These strategies can improve solubility by finding optimal (re-)folding conditions, but may also lead to activity loss of a protein (Singh & Panda, 2005). Further, when multiple enzymes in a pathway are insoluble, a single strategy may not be applicable. It is therefore desirable to select high-solubility enzymes to implement synthesis pathways.

To identify profitable synthesis pathway designs for experimental implementation, we couple in this paper the two synthesis tasks: *pathway construction* and *pathway implementation*. Using sequences from *UniProtKB* database (UniProt Consortium, 2017), we assemble a Protein Solubility Database (*ProSol DB*), that comprises confidence scores for the likelihood of an enzyme being soluble in *E. coli* and also allows for quick lookup using Enzyme Commission (EC) numbers. We used *ccSOL omics* (Agostini, Cirillo, Livi, Ponti, & Tartaglia, 2014) to compute solubility propensity scores for various proteins from *UniProtKB* in *E. coli.* Our database stores solubility confidence scores for 240,016 sequences, of which 34,046 sequences are associated with commonly used organisms. We develop a new workflow, termed Probabilistic Pathway Assembly with Solubility Scores (*ProPASS*), to couple *ProPath* (Yousofshahi et al., 2011), a method for constructing synthesis pathways, and *ProSol DB*. The workflow consists of first identifying synthesis pathways from a host to a target metabolite. Pathways are then ranked based on their length and yield. This ranking allows speedy identification of short length and high yielding pathways, which are desirable for experimental validation. *ProPASS* then recommends sequences based on their solubility confidence scores in *ProSol DB* to implement reactions along each pathway. We apply our workflow to identify implementations of synthesis pathways for seven target molecules. We analyze three test cases in detail. In all three cases, we show that *ProPASS* identifies one or more implementation that are predicted soluble. We further show that implementations published in the literature are recommended by *ProPASS* if catalogued in *UniProtKB*. As far as we know, this is the first description of a systematic workflow or method that explores using gene-specific sequence information to systematically identify host-compatible enzymes that can be used to implement synthesis pathways.

## Methods

### Assembling the Protein Solubility Database (*ProSol DB*)

To create a database of solubility confidence scores for various enzymes, we utilize annotated sequences in the *UniProtKB* database (UniProt Consortium, 2017). Protein sequences that have been either experimentally validated or manually analyzed are considered “reviewed” within the *UniProtKB/Swiss-Prot* database. While *UniProtKB/Swiss-Prot* currently contains over 550,000 sequences, only 240,016 sequences are associated with 4,652 Enzyme Commission (EC) numbers. *ProSol DB* is designed such that the EC numbers serve as key for database lookups, and the returned value contains the *UniProtKB/Swiss-Prot* sequence IDs, solubility confidence scores, and associated organisms. Solubility confidence scores are computed only once, using *ccSOL omics*, and stored locally in the database, eliminating the need to predict scores repeatedly, thus speeding up enzyme selection. *ccSOL omics* calculates solubility propensity of fragments within the sequences and utilizes neural networks to estimate confidence in solubility based on Fourier transform coefficients of the sequences (Agostini et al., 2014; Tartaglia, Cavalli, & Vendruscolo, 2007; Tartaglia, Pechmann, Dobson, & Vendruscolo, 2009).

While several solubility prediction tools are available, we calculate solubility confidence scores using *ccSOL omics*. The prediction accuracy of *ccSOL omics* reported is high (74%) for three independent datasets (Agostini et al., 2014). Further, the availability of the *ccSOL* omics code without a web interface facilitated the direct construction of our database. Several recent papers (Chang, Song, Tey, & Ramanan, 2013; Habibi, Hashim, Norouzi, & Samian, 2014; Khurana et al., 2018), provide assessment of the accuracy of various tools; however, each tool is trained on a different dataset. As our goal is to predict solubility of enzyme sequences from a broad range of organisms reported in *UniProtKB/Swiss-Prot*, we identified a subset of sequences in *UniProtKB/Swiss-Prot* for which experimental solubility labels are readily available. The labels were collected by comparing sequences from *UniProtKB/Swiss-Prot* to those in Target Track DB (Chen, Oughtred, Berman, & Westbrook, 2004). The total number of sequences in our independent dataset, referred to as *“the enzyme solubility test set”,* consisted of 716 sequences, where 462 are soluble and 254 insoluble. For the enzyme solubility test set, the accuracy of *ccSOL omics* was 73.46% using a 30% confidence threshold, compared to 46% for DeepSol S1, a recent tool for predicting solubility (Khurana et al., 2018). **Supplementary File 6** provides detailed accuracy results at various confidence thresholds for *ccSOL omics* and for DeepSol S1. While relatively small, the enzyme solubility test set is the most representative sample of *UniProtKB/Swiss-Prot*. The high prediction accuracy using *ccSOL omics* justifies its use when computing solubility for *ProSol DB*. Further, based on the analysis of the enzyme solubility test, we suggest using a solubility confidence score equal to or greater than 30% as “high confidence of solubility”.

### *ProPASS*: Coupling probabilistic pathway construction with protein solubility prediction

With the ability to lookup protein solubility confidence scores in *ProSol DB* based on EC numbers, we developed *ProPASS*, a workflow that couples pathway construction with identification of organism-specific enzymes for each step. The workflow (**Figure 1**) consists of a design step followed by an implementation step. Additional details can be found in **Supplementary File 1**.

**Figure 1:**
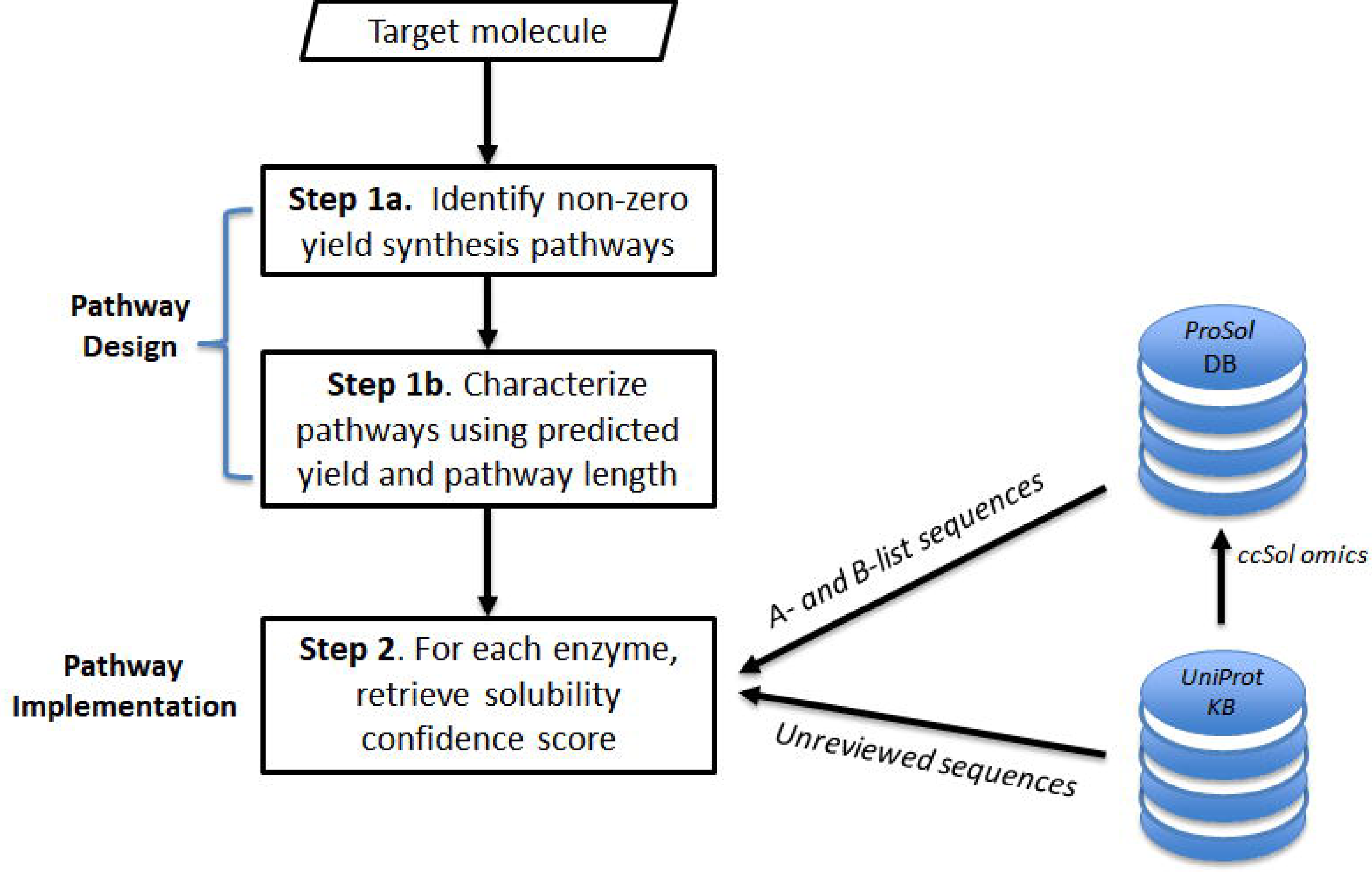
An overview of the steps used by *ProPASS* to identify synthesis pathways and their enzymatic implementations for a given target molecule. Solubility confidence scores from *ProSol DB* are retrieved for all sequences associated with a specific enzyme from the A-list or B-list organisms or computed on-the-fly using *ccSOL omics* for non-reviewed sequences in *UniProtKB*.

### Step 1 Synthesis pathway design

#### Step1a. Synthesis pathway identification

Given a target molecule, *ProPath* synthesizes pathways. *ProPath,* a graph-based probabilistic search algorithm, probabilistically selects synthesis pathways, from a target molecule terminating at a molecule within the host, using all reactions extracted from the KEGG database. We selected *ProPath* for pathway construction as it is computationally efficient and was shown effective in generating synthesis pathways with yield distributions similar to those obtained using limited-in-depth exhaustive search. Additionally, *ProPath* reproduces validated pathways published in the literature.

#### Step 1b. Synthesis pathway ranking

Identified pathways are ranked by two metrics: pathway yield and pathway length. These metrics are reported to users to allow the examination of different designs and their tradeoffs when identifying suitable pathway implementations for experimenters.

### Step 2 Synthesis pathway implementation

Given a synthesis pathway, the EC number for each of its enzymatic reaction is utilized to look up the associated *UniProtKB/Swiss-Prot* protein IDs in *ProSol DB*. The retrieved data also contains solubility confidence scores and organisms associated with each protein ID. The scores provide a *confidence level* that helps determine if a protein is soluble in *E. coli*. The scores should not be taken as any measure of relative solubility of a given enzyme in *E. coli*. In the case of non-matching protein IDs in *ProSol DB*, we allow the option to retrieve non-reviewed sequences from *UniProtKB*, and solubility confidence score for each sequence using *ccSOL omics* on ad hoc manner. Each reaction will have one or more recommended implementation sequence, along with its solubility confidence score and source organism.

## Results

### Overview of Protein Solubility Database (*ProSol DB*)

*ProSol DB* is considered the primary source of solubility confidence scores for *ProPASS* and consists of 240,016 sequences. *ProSol DB* is built using all reviewed protein sequences in *UniProtKB/Swiss-Prot*. The protein sequences in *ProSol DB* are associated with 4,652 EC numbers out of 6,896 EC numbers reported in KEGG (**Figure 2a**). We divide protein sequences organisms into two sets. The first set consists of 20 commonly used and well-studied organisms (e.g. *E. coli, S. cerevisiae, B. subtilis*, etc.) from which protein sequences can be easily acquired for experimental validation (A-list) (**Supplementary File 2**). The second set, the B-list, contains all other organisms associated with the remaining protein sequences associated with EC numbers. Protein sequences associated with the A-list organisms cover 2,427, or 52%, of all unique EC numbers available in *ProSol DB*, and 13% of all sequences in the database even though it consists of only 0.33% of the organismal diversity (**Figure 2b**). Example of solubility confidence scores present in our database can be found in **Table I**. The first column lists the EC number. The second column shows the associated *UniProtKB/Swiss-Prot* IDs for protein sequences associated with the EC number in the first column. The sequences are identified by their *UniProtKB/Swiss-Prot* IDs to make it easier to obtain any information related to these sequences from *UniProtKB/Swiss-Prot*. The third column contains a solubility confidence score for each protein sequence. The fourth and fifth columns show organisms associated with each protein sequence. When an EC number is queried, a list of protein IDs associated with the A-list organisms is returned. If none are associated with the A-list, a list of protein IDs associated with the B-list is returned. Further examples of solubility confidence scores within *ProSol DB* can be found in **Supplementary File 3.**

**Figure 2:**
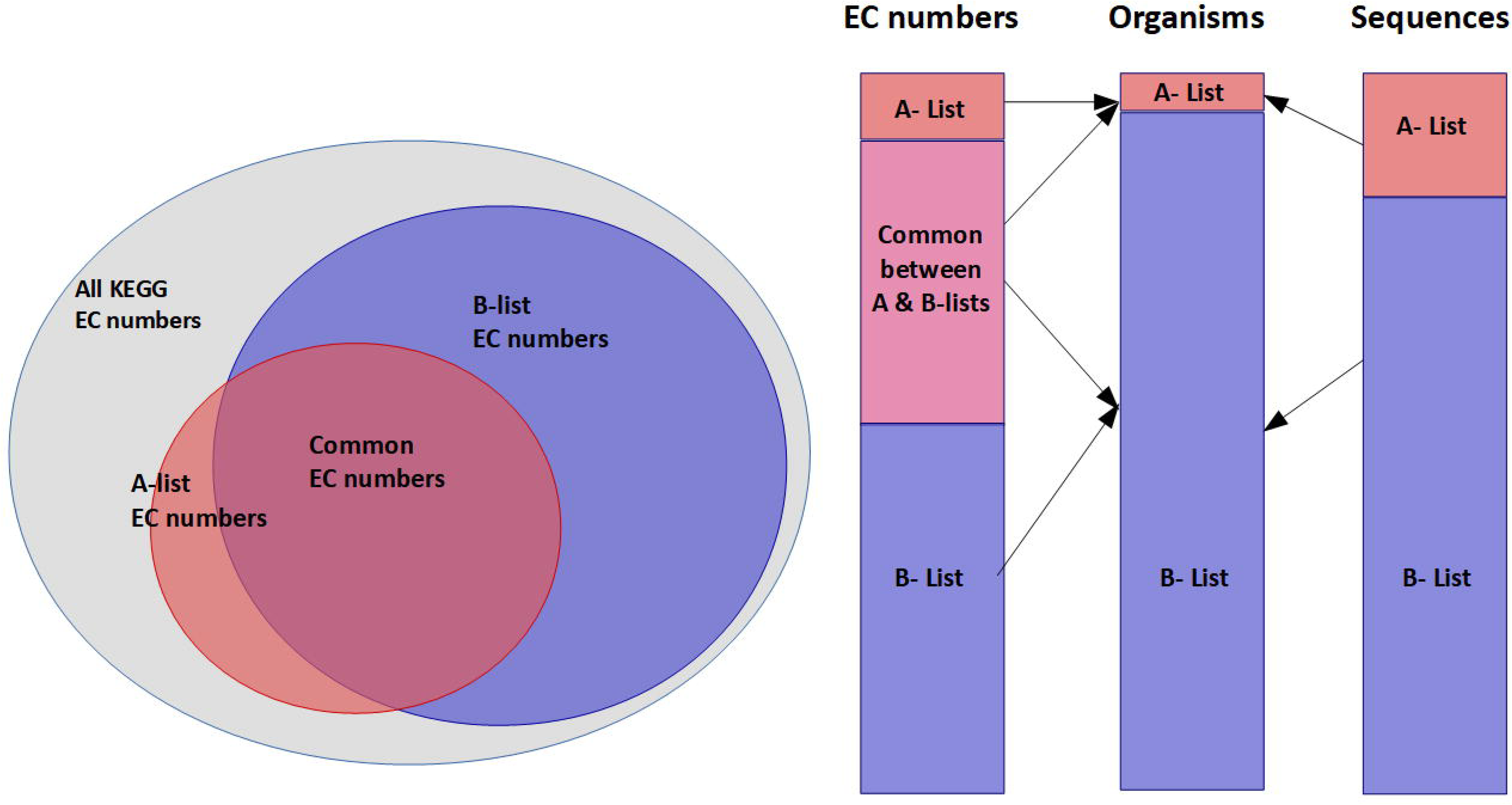
Statistics related to *ProSol* DB. (a) Venn diagram showing sets of all EC numbers in KEGG (6,896), those covered by the A-list (2,427), and those covered by the B-list (4,350). The intersection between the A-list and the B-list consists of 2,125 EC numbers covering 46% of EC numbers available in *ProSol* DB. (b) Each stack shows information related to the A-list and B-list organisms. The first bar shows tally of EC numbers in A-list, B-list, and their intersection. The second bar shows that the A-list consists of 20 organisms, which is only 0.33% of the organism diversity yet covers 52% of all the EC numbers in the database. The third bar shows that 13% of all protein sequences in *ProSol DB* are covered by the A-list organisms. These statistics demonstrate that the number of verified enzyme sequences and EC numbers are overrepresented in the A-list, which comprises many model organisms.

We analyzed correlations of confidence scores across EC classes and sourcing organisms. If such correlations existed, they could guide pathway synthesis and implementation algorithms by favoring EC’s or organisms with increased propensity for soluble proteins. Since *ProSol DB* is the largest database of solubility confidence scores in *E. coli*, we used it as a resource to investigate any potential correlation between propensity for soluble expression and EC numbers (**Figure 3**). For this analysis, we only considered EC numbers that are associated with 10 or more protein sequences in *ProSol DB*. The average predicted solubility confidence score for the 1,600 enzymes that meet this criterion is 54.3 (**Figure 3a**). For each EC number, the range of solubility confidence scores (solubility range) varied tremendously (**Figure 3b**), with over 96% of ECs having scores that varied by a difference of 75 or more. Analyzing the standard deviation of average solubility confidence score per EC number shows that they are also spread over similar wide ranges (**Figure 3c**). There was low correlation between the score ranges and average solubility confidence scores (**Figure 3d**) as analyzed using Spearman’s Rank Test (Spearman’s rank coefficient = 0.00897)

**Figure 3:**
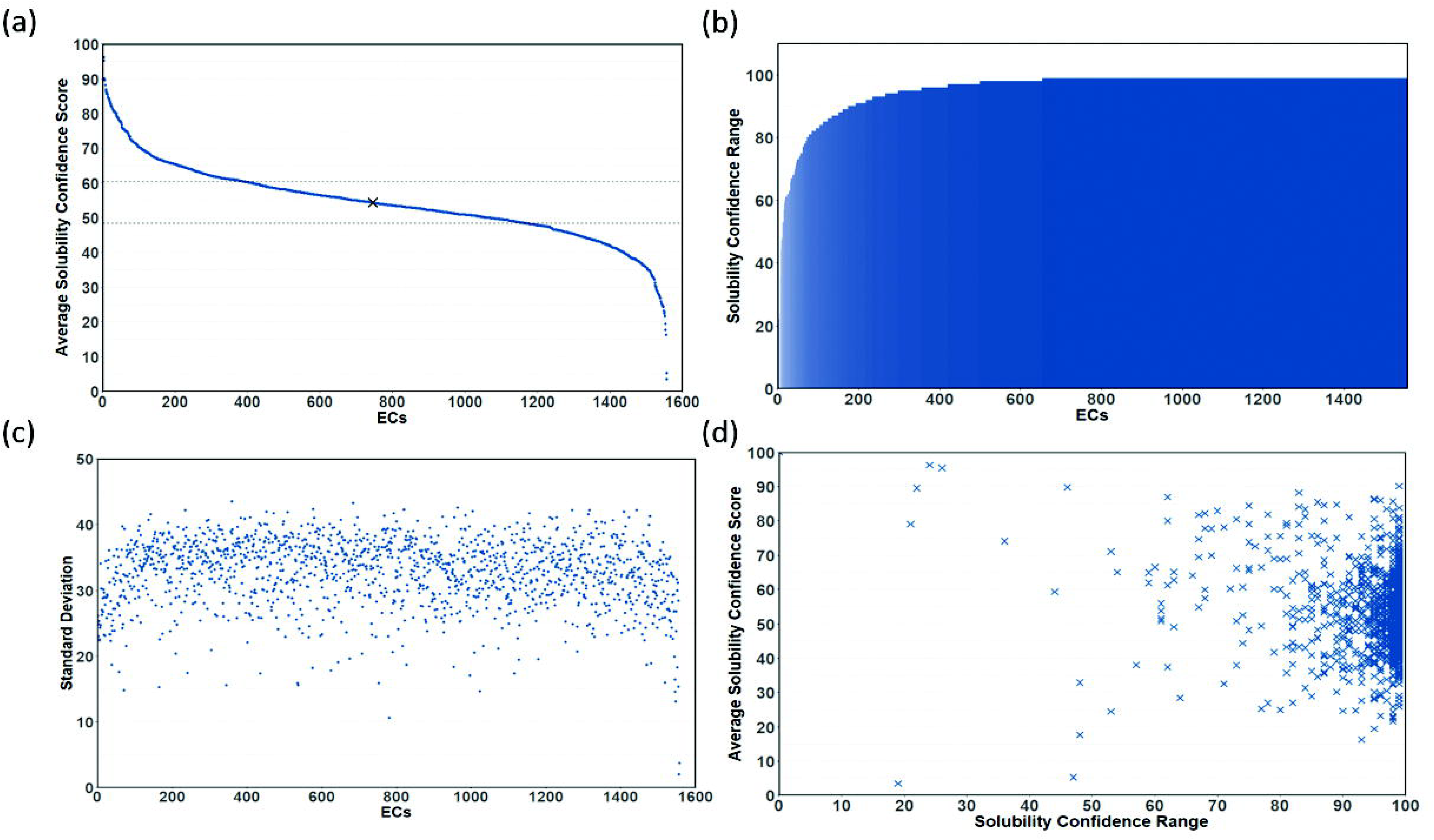
Correlation analysis between solubility confidence scores and EC numbers for enzymes with 10 or more sequences in *ProSol DB*. (a) Average solubility confidence scores for enzymes, sorted from highest to lowest average score. (b) Predicted solubility confidence score ranges for each enzyme, sorted from smallest to largest. (c) Standard deviation of solubility confidence scores for each enzyme, sorted as in panel (a). (d) Scatter plot of solubility confidence score ranges and average scores. There is low correlation between the score ranges and average solubility confidence scores as determined using Spearman’s Rank Test (Spearman’s rank coefficient = 0.00897).

Further, we investigated the correlation between solubility confidence scores and the similarity of the source organisms to *E. coli* (**Figure 4**). The phylogenetic distance from *E. coli* was computed using phyloT, a tool for the annotation and visualization of phylogenetic trees (Letunic & Bork, 2016). We only considered organisms that had at least 100 solubility confidence scores in *ProSol DB*. The distances varied between 0 and 40 (**Figure 4a**), where a score of 0 indicated a specific strain of *E. coli*, and a score of 40 indicated a very distant species such as *Drosophila melanogaster* (fruit fly). The average solubility confidence score per organism was 53.43 (**Figure 4b**). The correlation between the phylogenetic distance and average solubility confidence score per organism (**Figure 4c**) showed low positive correlation as calculated using Spearman’s Rank Test (Spearman’s rank coefficient = 0.176919). We conclude from **Figures 3** and **4** that there is low to no correlation between propensity for soluble expression and EC numbers, and between solubility confidence scores and the similarity of the source organisms to *E. coli*. Therefore, when implementing a pathway, protein sequences are selected with respect to their individual solubility confidence scores.

**Figure 4:**
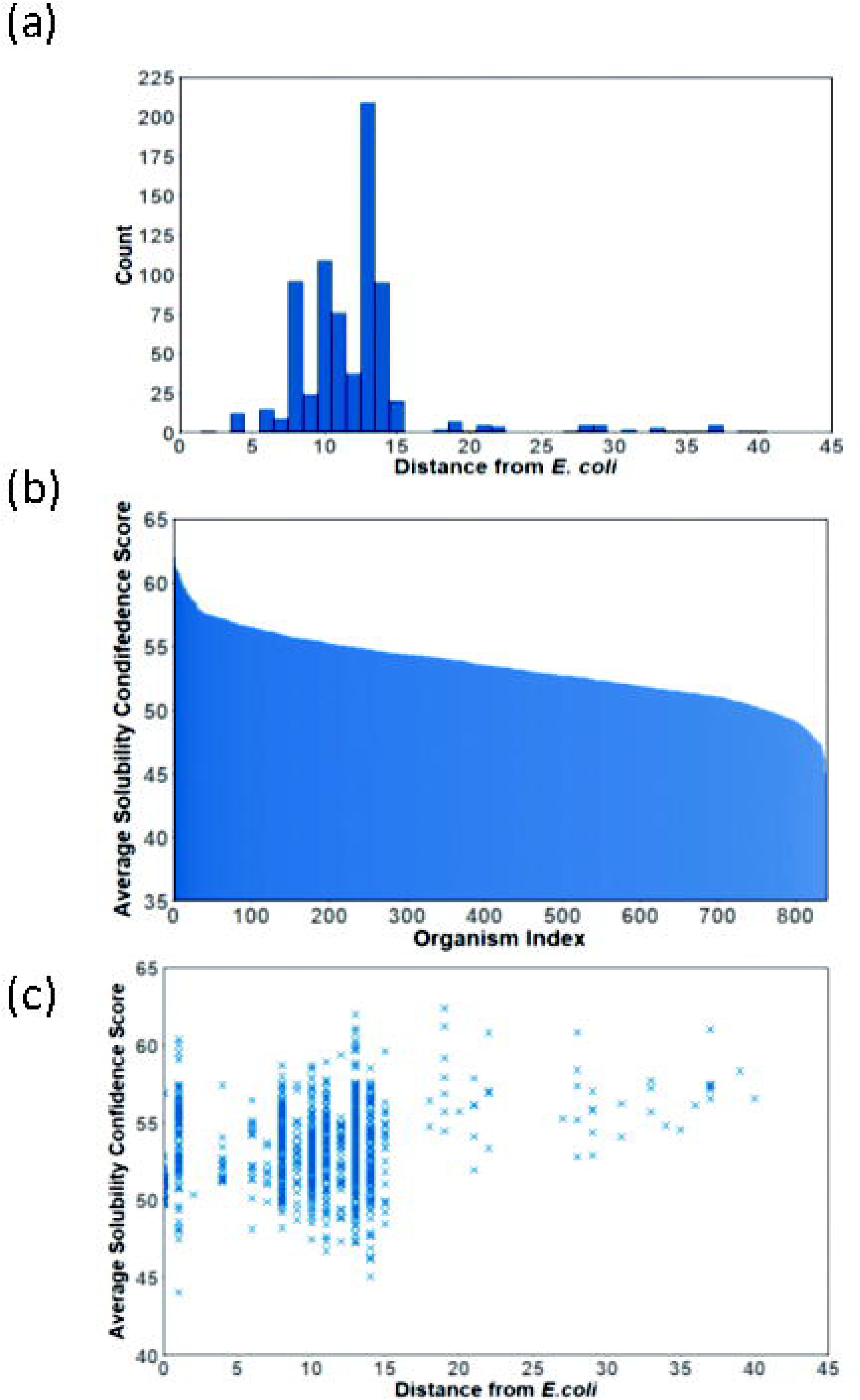
Correlation analysis between predicted solubility scores and sourcing organisms for those with 100 or more sequences in *ProSol* DB. (a) Histogram of phylogenetic distances from *E. coli.* (b) Average predicted solubility per organism, ranked from largest to smallest predicted solubility values. (c) Scatter plot of average predicted solubility per organism vs. phylogenetic distance from *E. coli* indicates low positive correlation as calculated using Spearman’s Rank Test (Spearman’s rank coefficient = 0.176919).

### Validating *ProPASS* workflow

We used the *ProPASS* workflow to identify synthesis pathways and their potential implementation for several commercially useful biomolecules. The targets include industrial precursors used in the production of plastics, printing ink, pharmaceuticals (2,3-butanediol (Cho et al., 2015), *cis*-muconic acid (Weber et al., 2012)), biofuels (triacylglycerols (Radakovits et al., 2010) and fatty acid methyl esters (Nawabi et al., 2011)), a commodity chemical that is widely used in industry (3-hydroxypropanoic acid (Della Pina, Falletta, & Rossi, 2011)) and biological precursors (isopentenyl diphosphate (Kim & Keasling, 2001) and *myo*-inositol (Shiue & Prather, 2014)). The test cases show that 20% − 100% of the identified pathways have solubility confidence scores for enzymes catalyzing all their reactions from the reviewed sequences present in *ProSol DB* (**Table II**). For example, for target molecule *myo*-inositol, *ProPASS* identified 384 pathways, of which 235 pathways (61% of the identified pathways) had solubility confidence scores for all reactions. The coverage is expected to further increase if the user wishes to utilize unreviewed sequences. Three of the seven test cases mentioned above were analyzed in detail of which two are presented below (3-hydroxypropanoic acid and 2,3-butanediol), and one in **Supplementary Files 4 and 5** (*cis*-muconic acid). In all three test cases, we constructed the pathway using *ProPath* and ranked the designs based on yield and length. The identified pathways were ranked based on the two design criteria: pathway yield and pathway length. Then, for the 5-top yielding and for the 5-top shortest pathways, we used *ProPASS* to identify potential implementations based on solubility confidence scores from *ProSol DB*. We compared the recommended implementations with published implementation of pathways reported in the literature.

### 3-Hydroxypropanoic acid

We applied *ProPASS* to synthesize 3-hydroxypropanoic acid (3-HP). Since 3-HP production through the *pdu* pathway is the only known pathway from glycerol, we only explored those originating from glucose and identified 112 synthesis pathways (**Figure 5**). Pathways lengths varied from 2 to 14 and yield values varied from 0.271 to 0.41 mmol/mmol of glucose/g CDW/h.

**Figure 5:**
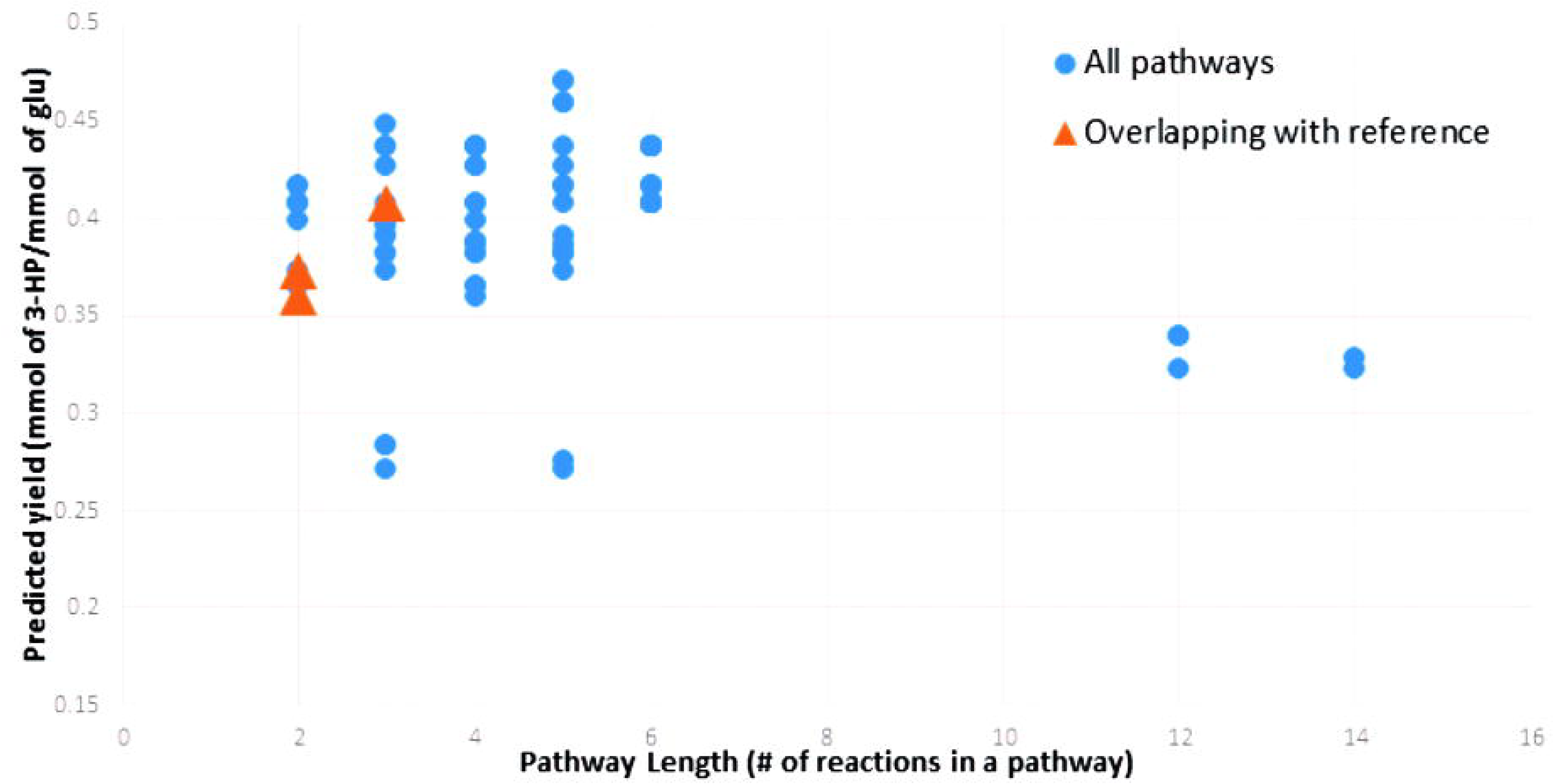
Predicted yields and corresponding lengths for pathways producing 3-hydroxypropanoic acid identified by *ProPASS*. Each point on the plot represents one pathway. The three orange triangles coincide with pathways reported in the literature.

When ranked based on yield (**Table IIIa**), the top pathway had five steps and yield of 0.408 mmol/mmol of glucose/g CDW/h. However, two of the five reactions had no solubility data in *ProSol DB.* Other pathways in the five top-ranked pathways had three or five steps. Two of the three-step pathways, yielding a value of 0.407 mmol/mmol of glucose/g CDW/h, had complete solubility confidence scores.

When ranked based on length (**Table IIIb**), the top five had only 2 reaction steps – from malonate semialdehyde to 3-HP. All five pathways had similar yields (0.365 – 0.373 mmol/mmol of glucose/g CDW/h). Only two of the give pathways had complete solubility data.

We compared the predicted solubility confidence scores of the enzymes used in the published literature to the highest solubility confidence score alternatives identified by *ProPASS* (**Figure 6**). Of the identified pathways, three have been experimentally used previously for 3-HP production either through the propionyl-CoA pathway (**Figure 6a**) (Luo et al., 2016), the malonyl-CoA pathway (**Figure 6b**) (Cheng et al., 2016), or β-alanine pathway (**Figure 6c**) (Song et al., 2016). In the case of the enzyme malonyl-CoA reductase in the malonyl-CoA pathway (**Figure 6b**), *ProPASS* identified a solubility confidence score of 74, while the corresponding enzyme from Cheng et al. (Cheng et al., 2016) has a score of 97. *UniProtKB* did not catalogue the sequences used by Cheng et al., which precluded their enzyme from the *ProPASS* workflow results. For 3-HP synthesis through the propionyl-CoA and the β-alanine pathways, our workflow identified sequences with high or similar solubility confidence scores when compared with sequences used in the literature.

**Figure 6:**
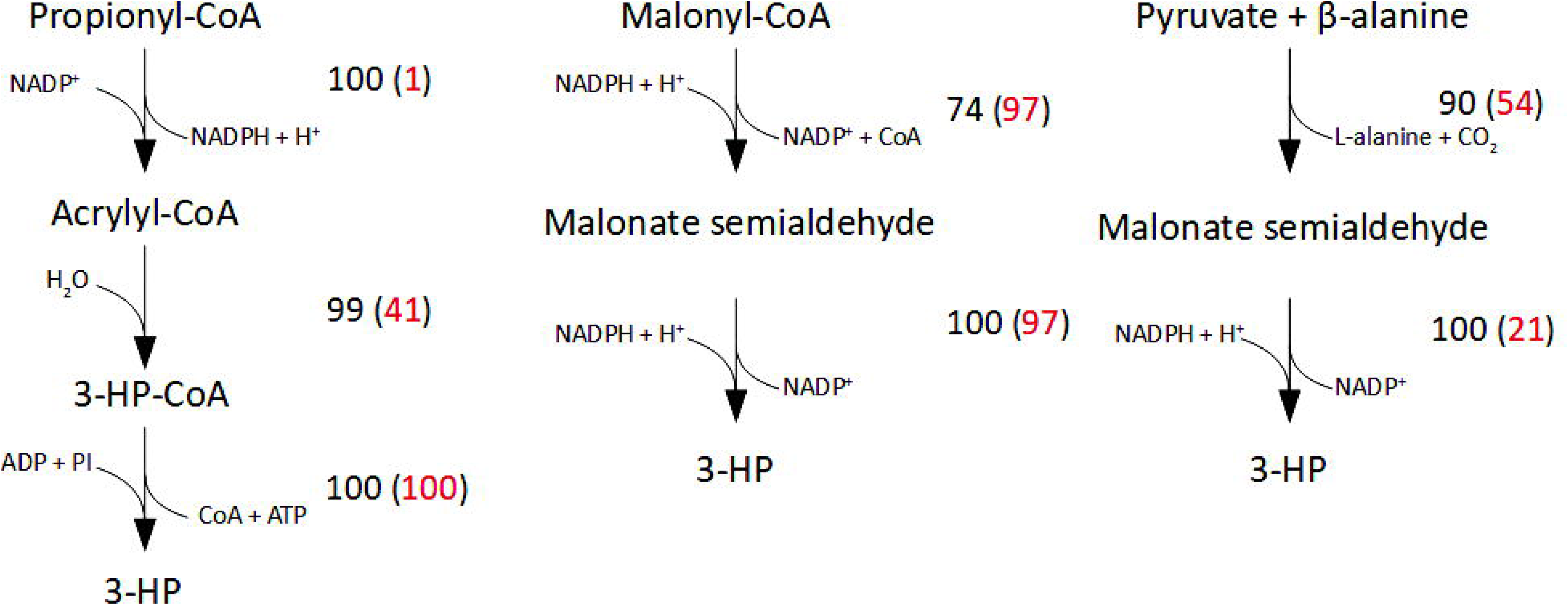
Three pathways for producing 3-hydroxypropanoic acid (3-HP) in *E. coli* were identified by *ProPASS* and proposed by (a)Luo et al. (2016), (b)Cheng et al. (2016), and (c)Song et al. (2016). Two predicted solubility scores are provided for each enzymatic reaction. The black score is for a sequence from organism recommended by *ProPASS* and the red score is for the corresponding sequence that was used experimentally.

### 2,3-Butanediol

*ProPASS* identified only one 2-step synthesis pathway that produces 2,3-butanediol using the (*R,R*)-butanediol dehydrogenase and acetolactate decarboxylase enzymes (**Table IV**). The highest solubility confidence scores for sequences identified by *ProPASS* from the A-list organisms were 87 (*S. cerevisiae*) and 83 (*L. lactis*), respectively. Expanding the search to include sequences from the B-list did not yield higher scores. Upon expanding the search to include reviewed and non-reviewed sequences from *UniProtKB*, *ProPASS* identified sequences with solubility confidence scores of 100 (*L. lactis*) for (*R,R*)-butanediol dehydrogenase and 99 (*E. aerogenes*) for acetolactate decarboxylase enzymes. We compared our findings with those in the literature and found that this 2-step pathway is the only known synthesis pathway for 2,3-butanediol. Further, the selected enzymes match an experimentally validated pathway by Yan et al. (Yan, Lee, & Liao, 2009), where enzyme sequences were selected from two A-list organisms, *B. subtilis* and *Klebsiella pneumonia.* Both sequences have high-predicted solubility scores, comparable to those identified in the non-reviewed sequences. As these sequences were not catalogued in *UnitProtKB/Swiss-Prot*, *ProPASS* did not identify them as possible sequence options.

## Discussion

Productive integration of heterologous pathways into model host organisms is vital for metabolic engineering. Key design steps where computational tools can significantly expedite the engineering cycle are *pathway construction* to identify reactions required to connect the host to a target metabolite, and *parts selection* to choose specific biochemical parts (such as enzymes, promoters, etc.) to express in the host. While current pathway construction tools can identify high-yielding pathways, they cannot be directly translated into experiments without the parts selection step. Next-generation synthesis tools must provide detailed biochemical and biophysical information about the biological parts that can be directly translated and integrated into the engineering design-build-test-learn cycle. Such information includes, but is not limited to, the sequence of heterologous enzymes, their activity, solubility, stability, etc.

Aiming to create a next-generation synthesis tool, we expanded in this work the scope of conventional pathway construction to identify potential sources from which each of the pathway enzymes can be isolated using protein solubility as a criterion of parts selection and host-compatibility. A specific amino acid sequence of enzymes in pathway determines many outcomes such as kinetics, stability, and actual yield. Solubility however is a barrier in achieving these outcomes. When a recombinant enzyme is expressed as an insoluble aggregate (known as inclusion bodies), they retain none or a fraction of the catalytic activity, except in a few cases (Choi et al., 2011; Lin, Zhou, Wu, Xing, & Zhao, 2013; Nahalka & Nidetzky, 2007; Zhou, Xing, Wu, Zhang, & Lin, 2012). An expressed recombinant enzyme therefore must be soluble in the heterologous host to ensure product yield and titer.

Current metabolic engineering practices utilize an ad hoc approach to enzyme sourcing based on domain knowledge or practical considerations such as accessibility to organisms from which enzymes can be sourced. For example, *n*-butanol synthesis pathway is often sourced entirely from natural producers, *Clostridia* (Inui et al., 2008; Nielsen et al., 2009). However, when solubility and expression of heterologous enzymes (i.e., host compatibility) are taken into consideration, several groups have found that a “mix-and-match” approach is often more profitable than sourcing all enzymes from a single organism, as was demonstrated when implementing the *n*-butanol pathway in yeast (Eric J Steen et al., 2008) and *E. coli* (Bond-Watts, Bellerose, & Chang, 2011). More drastic examples of such sourcing paradigm were demonstrated in by both Keasling and Smolke groups when synthesizing complex natural products (Galanie et al., 2015; Ro et al., 2006). Assembly of such “mix-and-match” pathways has been significantly aided by decreasing costs of gene synthesis. However, such empowerment also leads to choice overload. We posit that host-compatibility in the form of solubility can provide a powerful guide to prune down the options to most profitable choices. Enzyme engineering, for improved catalytic efficiency, substrate and product preferences, and solubility, is traditionally used as a strategy to overcome issues with product synthesis. It can, however, be time-consuming, even when aided by computational protein engineering tools (e.g. (Hassanpour, Ullah, Yousofshahi, Nair, & Hassoun, 2017; Sachsenhauser & Bardwell, 2018)), especially if several enzymes within a pathway are insoluble. Thus, implementing pathways with knowledge that the enzymes selected have a high propensity for solubility in the host can reduce the traditional trial-and-error approach to enzyme selection. The product of *ProPASS* is a listing of synthesis pathways and recommended implementations. *ProPASS* therefore is a tool to aid in exploring the design space of synthesis pathways and their implementation.

Therefore, protein solubility is used as a rating and/or selection criterion before the implementation phase. In most cases, multiple implementation options based on solubility confidence scores are available to choose from. Since solubility confidence scores do not affect maximum theoretical yield, they do not provide a ranking criterion for pathway design. Rather, they provide a guideline for the implementation phase. Thus, *ProPASS* advances the state of pathway synthesis by exploring the underlying biophysical design space, leading to more profitable design implementation options when compared to using pathway synthesis tools that only explore the abstract metabolic space. Such incorporation of sequence information is distinct from previous descriptions where protein sequence and structure information has been used to inform putative promiscuous enzymatic activity to assemble non-natural synthetic pathways (Brunk, Neri, Tavernelli, Hatzimanikatis, & Rothlisberger, 2012; Erb, Jones, & Bar-Even, 2017).

To allow for enzyme selection, we created a database of predicted solubility confidence scores, *ProSol DB*, for reviewed protein sequences from the *UniProtKB/Swiss-Prot* database. *ProSol DB* contains 240,016 sequences and links EC numbers with protein sequences, their source organism, and their solubility confidence scores in *E. coli*. The solubility confidence scores were calculated using *ccSOL omics*, with reported accuracy comparable with other available solubility prediction tools. Future improvements in solubility prediction algorithms can be used to update *ProSol DB*, thus providing more confidence regarding enzyme selection. Analysis of solubility confidence scores and enzymes revealed that sequences associated with a particular enzyme have a wide range of associated scores. Further analysis showed no correlation between phylogenetic distance from *E. coli* and solubility confidence scores. Our findings emphasize that solubility is a property of the encoding sequence, and not a function of the host organism or associated EC number. As far as we know, *ProSol DB* is the largest database of enzyme solubility confidence scores for heterologous enzymes in *E. coli* and serves as a resource for pathway synthesis tools as well as for meta-analysis of sequence independent factors that affect solubility of enzymes in *E. coli*.

We validated the *ProPASS* workflow by comparing predicted pathway implementations with those published in literature for three target molecules, 3-hydroxypropanoic acid, 2,3-butanediol, and *cis*-muconic acid. In all cases, we identified a significant number of pathway implementations for which solubility confidence scores were available through *ProSol DB*. The availability of data was dependent on the studied target molecule. In the case of *myo*-inositol, 61% of identified pathways had confidence scores for each enzyme, while in the case of *cis-*muconic acid only 20% of identified pathways had such solubility data. Several observations can be drawn based on examining the three detailed test cases. When considering solubility, longer pathways have a higher chance of having enzymes with low solubility confidence scores. For example, the *cis-* muconic acid synthesis pathway with the highest yield has a pathway length of 10, with one of the steps having a low solubility confidence score of 22. In contrast, the pathway with the highest confidence of having a soluble enzyme for each step is a three-step pathway but has only half the yield of the highest-yield pathway. *ProPASS* can provide several implementation alternatives, from which one or more pathways can be selected for experimental testing.

When compared to solubility confidence scores of published implementations for the three test cases, *ProPASS* recommended the published implementations provided they were in *UnitProtKB.* In some cases, *ccSOL omics* predicted the published successful enzyme implementation as insoluble, with solubility confidence scores less than 30. This was the case for three (e.g., the first step for producing 3-HP from propionyl-CoA, **Figure 6a**, the second step of producing 3-HP from pyruvate and β-alanine, **Figure 6c**, and the first step of producing *cis-*muconic acid from *para*- hydroxybenzoic acid, **Figure S2b**) of twelve comparisons with known soluble enzymes reported in the literature (**Figure 6, Table IV,** and **Figure S2**). Such mispredictions are expected considering that the accuracy of *ccSOL omics* for the enzyme solubility test set was evaluated to be 73.46%. When the solubility confidence score for a given enzyme in a given pathways was low, it was possible to utilize non-reviewed sequences in *UniProtKB* to identify higher score alternatives. This was the case when synthesizing 2,3-butandiol. As more reviewed sequences are added to *UniProtKB/Swiss-Prot,* the reliance on non-reviewed sequences will diminish. Importantly, *ProPASS* can provide alternative implementations that would not have been easily discoverable otherwise. All test cases discussed here are limited to using *E. coli* as host. However, development of solubility prediction tools for other organisms would allow extending *ProPASS* to other hosts.

Strain optimization, the process by which opportunities for metabolic re-direction are identified to ensure maximal target formation, is a crucial step that ensures the profitable integration of heterologous synthesis pathways into model host organisms. While several computational tools (e.g., (Burgard, Pharkya, & Maranas, 2003), (Patil, Rocha, Förster, & Nielsen, 2005), and (Ranganathan, Suthers, & Maranas, 2010)) were proposed to identify the most profitable modifications, almost none take into account implementation concerns during optimization. One exception is *CCOpt* (Yousofshahi, Orshansky, Lee, & Hassoun, 2013), which addresses the uncertainty in precise tuning of enzyme levels and imposes probabilistic flux capacity constraints that capture the uncertainty in tuning enzyme levels during implementation. In this work, we treated the two design steps (pathway synthesis and strain optimization) as independent. A non-competitive (in terms of yield) synthesis pathway during the pathway synthesis step may provide higher yields than other synthesis options once strain optimization is performed. When developed judiciously to limit the design space, co-optimization tools that are aware of design choices associated with the underlying implementation can potentially yield more profitable experimental guidance.

## Acknowledgements

We thank Emily Chicklis, an REU student, for her help with some figure illustrations.

